# Development of ONT-cappable-seq to unravel the transcriptional landscape of *Pseudomonas* phages

**DOI:** 10.1101/2022.03.18.484859

**Authors:** Leena Putzeys, Maarten Boon, Eveline-Marie Lammens, Konstantin Kuznedelov, Konstantin Severinov, Rob Lavigne

**Affiliations:** Department of Biosystems, Laboratory of Gene Technology, KU Leuven, Leuven 3001, Belgium; Waksman Institute, Rutgers, The State University, Piscataway, NJ 08854, USA

## Abstract

RNA sequencing has become the method of choice to study the transcriptional landscape of phage-infected bacteria. However, short-read RNA sequencing approaches generally fail to capture the primary 5’ and 3’ boundaries of transcripts, confounding the discovery of key transcription initiation and termination events as well as operon architectures. Yet, the elucidation of these elements is crucial for the understanding of the strategy of transcription regulation during the infection process, which is currently lacking beyond a handful of model phages. To this end, we developed ONT-cappable-seq, a specialized long-read RNA sequencing technique that allows end-to-end sequencing of primary prokaryotic transcripts using the Nanopore sequencing platform. We applied ONT-cappable-seq to study transcription of *Pseudomonas aeruginosa* phage LUZ7, obtaining a comprehensive genome-wide map of viral transcription start sites, terminators, and complex operon structures that fine-regulate gene expression. Our work provides new insights in the RNA biology of a non-model phage, unveiling distinct promoter architectures, putative small non-coding viral RNAs, and the prominent regulatory role of terminators during infection. The robust workflow presented here offers a framework to obtain a global, yet fine-grained view of phage transcription and paves the way for standardized, in depth transcription studies for microbial viruses or bacteria in general.

## INTRODUCTION

Over the last decade, high-throughput sequencing approaches have fuelled a new era of molecular research in bacteria and their viral predators, bacteriophages. The surge in microbial transcriptome studies has provided major insights in the fundamentals of their transcription strategies and mechanisms of gene regulation (1–3). To date, short-read RNA sequencing (RNA-seq) technology remains the leading platform to record the global transcriptional state of phage-infected bacteria (4– 7). However, standard RNA-seq techniques often fail to capture key transcriptional events such as transcription initiation and termination, as these approaches do not discriminate between primary and processed transcripts. As a consequence, the annotation of transcription start sites (TSS) and transcription termination sites (TTS), representing the original boundaries of primary transcripts, is generally confounded by the extensive *in vivo* processing of transcripts in the RNA pool (1). Prior to sequencing, various strategies can be used to enrich the fraction of primary transcripts in the RNA population and facilitate global TSS determination in prokaryotes. The main strategy is differential RNA-seq (8), in which processed transcripts are depleted from the RNA pool using terminator exonuclease treatment. Conversely, Cappable-seq relies on a targeted enrichment of primary RNAs by enzymatically labelling their 5’ triphosphate group (9). These strategies have yielded comprehensive transcription initiation maps of numerous bacteria and have recently also been used to study viral infection (10, 11). However, as both techniques rely on short-read sequencing technology, termination sites remain more difficult to delineate and information on transcript continuity and operon complexity is lost during sample processing.

Recently, long-read sequencing technologies have entered the transcriptomics fields and provide the opportunity to sequence full-length transcripts, enabling more straightforward identification of transcriptional boundaries (12–14). For example, SMRT-Cappable-seq uses the PacBio SMRT (Single-molecule-real-time) platform to sequence full-length primary transcriptomes and reveal transcriptional landmarks and operon structures in bacteria (15). However, as this method uses size selection to enrich for longer transcripts, it is less suited for the exploration of phage transcriptional landscapes, which are often hallmarked by high gene densities and short open reading frames. In parallel, Oxford Nanopore’s long-read sequencing technology (ONT) is rapidly improving in quality and is emerging as a cost-effective technique in microbial transcriptomics (12, 16–18).

In this study, we present a nanopore-based Cappable-seq approach (ONT-cappable-seq), which can be especially useful for resolving regulatory features in densely coded viral genomes. The power of full-length primary transcriptome sequencing was just recently demonstrated for the eukaryotic severe acute respiratory syndrome coronavirus 2 (SARS-CoV-2), where it enabled a better understanding of its complex transcriptome (19). The ONT-cappable-seq method outlined in this work enables end-to-end sequencing of primary prokaryotic transcripts to unravel the complex transcriptomes of microbial viruses and their bacterial hosts.

As a proof-of-concept, we elucidated the transcriptional properties of *Pseudomonas aeruginosa* virus LUZ7. Like other relatives of coliphage N4, LUZ7 has a peculiar transcriptional strategy that relies successively on three different RNA polymerases (RNAP) (20, 21). Upon phage infection, a large virion-encapsulated RNAP (vRNAP) is co-injected with the viral DNA and initiates transcription of the early genes from single-stranded hairpin promoters. Among the early phage gene products, the heterodimeric RNAP (RNAPII) and single-stranded DNA-binding protein Drc activate transcription of the middle genes (22). Finally, the host RNAP takes over during the final infection stage and carries out transcription from late phage promoters, which weakly resemble the σ^70^ promoter consensus motif (23). However, no middle and late viral promoters have been definitively identified in LUZ7 and its closest relatives, despite the availability of RNA-seq data for one of these phages (24). This gap of knowledge and a complex transcriptional strategy makes LUZ7 an interesting model to benchmark our method and at the same time unravel the unknowns of LUZ7 genome transcription.

## MATERIALS AND METHODS

### 1. Bacteriophage propagation

*P. aeruginosa* strain US449 was cultured at 37°C in Lysogeny Broth (LB) medium. To amplify *Pseudomonas* phage LUZ7 (25), a culture of US449 was grown to an optical density at 600 nm (OD_600_) of 0.3 and infected with a high titre lysate of LUZ7, followed by overnight incubation at 37°C. Afterwards, the phage lysate was purified with polyethylene glycol 8000 (PEG8000) and stored in phage buffer at 4°C as described elsewhere (26).

### 2. Bacterial genome extraction, nanopore sequencing and hybrid assembly

The genome of *P. aeruginosa* strain US449 was previously sequenced using Illumina short-read sequencing technology (NCBI accession number GCF_001454415.1). To complete the reference genome, the short reads were complemented with long reads obtained after sequencing the genomic DNA of the bacteria using Nanopore sequencing technology. First, high-molecular weight bacterial genomic DNA was extracted using the DNeasy UltraClean Microbial Kit (Qiagen, Hilden, Germany). Next, the DNA sample was carefully prepared using the Rapid Barcoding Sequencing Kit (SQK-RBK004), loaded on a MinION flow cell (FLO-MIN 106, R9.4) and sequenced for 24-48h. The base-calling, demultiplexing and trimming of the raw nanopore reads was performed using Guppy (v3.4.4) and Porechop (v0.2.4) (https://github.com/rrwick/Porechop). The final quality of the sequencing run and the read lengths was evaluated using NanoPlot (v1.28.2) (27). Afterwards, both short and long-read sequencing datasets were integrated to resolve the US449 genome using *de novo* hybrid assembly. For this, the Unicycler tool (v0.4.8) was used with default process parameters, followed by an additional polishing step (28, 29). The resulting high-quality genomic sequence of *P. aeruginosa* US449 was annotated with Prokka (v1.14.6) using default settings (30). This assembly and annotation was used for further analysis of the transcriptome data. The resolved genome of *P. aeruginosa* strain US449 was deposited in NCBI GenBank (accession number CP091880).

### 3. Infection conditions and total RNA extraction

An overnight culture of *P. aeruginosa* US449 was inoculated in 20 mL fresh LB medium and grown to the early exponential phase (OD_600_ = 0.3). At this moment, a 4.5 mL culture sample was collected to serve as an uninfected control sample (t = 0 min). The remaining culture was infected with phage LUZ7 with a multiplicity of infection (MOI) of 50 to ensure a synchronous infection. The culture was incubated at 37°C and 4.5 mL samples for RNA isolation were collected at different timepoints during infection that represent the early (5 min), middle (10 min) and late (20 min) infection stage, as determined from previous growth experiments (21, 31). All collected samples were immediately mixed with 0.5 mL of stop mix solution (95% v/v ethanol, 5% v/v phenol) and snap-frozen in liquid nitrogen to preserve the transcriptional state of the cells. Prior to total RNA extraction, high infection rates (>95% cells infected) were confirmed by comparing the number of colony-forming units (CFU/mL) in the uninfected control and the 5 minutes post-infection sample after overnight incubation.

Next, the samples were thawed on ice and centrifuged (20 min, 4,000 *g*, 4°C). Total RNA was isolated from the cell pellet by lysozyme treatment, followed by hot phenol extraction and subsequent ethanol precipitation. The samples were treated with DNase I and cleaned using ethanol precipitation and subsequent RNA Clean & Concentrator-5 spin-column purification (Zymo Research). Successful removal of genomic DNA was confirmed by PCR using a host-specific primer pair (Supplementary Figure S1). The purity and concentration of the RNA samples were measured by the spectroscopic SimpliNano device (Biochrom US, Inc.) and a Qubit 4 fluorometer using an RNA HS assay kit (ThermoFisher Scientific), respectively. Evaluation of RNA integrity was performed using an Agilent 2100 Bioanalyzer system in combination with the RNA 6000 Pico Kit, and only samples with RNA integrity numbers (RIN) > 9 were used for downstream processing and sequencing.

### 4. In vitro transcription of RNA spike-in

The HiScribe T7 High Yield RNA Synthesis Kit (New England Biolabs) was used for the *in vitro* synthesis of RNA transcripts from a FLuc control template, according to manufacturer’s guidelines. The DNA template was removed by DNase I treatment after incubation at 37°C for 30 minutes. After ethanol precipitation and additional spin-column purification using the Zymo RNA Clean & Concentrator-5 kit, the quality and quantity of the purified transcripts was assessed using a SimpliNano spectrophotometer. The RNA products were run on a RNA 6000 Pico chip using a Agilent 2100 Bioanalyzer device to evaluate transcript integrity and length (1.8 kb).

### 5. ONT-cappable-seq library preparation and sequencing

#### Enrichment of primary transcripts

The enrichment of primary transcripts was performed by an adapted protocol based on the (SMRT)-Cappable-seq method (9, 15). Total RNA (5 µg) was supplemented with 1 ng of RNA spike-in in a total volume of 30 µL, incubated for 5 minutes at 65°C, and placed on ice. The RNA was capped by adding 0.5 mM of 3’-desthiobiotin-GTP (3’-DTB-GTP) (New England Biolabs), 50 units of Vaccinia Capping Enzyme (New England Biolabs), 5 µL of 10X VCE Buffer (New England Biolabs) and 0.5 units of yeast pyrophosphatase (New England Biolabs), followed by incubation for 40 minutes at 42°C. The capped RNA was cleaned by spin-column purification using the Zymo RNA Clean & Concentrator-5 kit with a total of four washes to ensure complete removal of unincorporated 3’DTB-GTP. Next, RNA molecules were polyadenylated by incubation for 15 minutes at 37°C in the presence of 20 units of *E. coli* Poly(A) Polymerase (New England Biolabs) and 1 mM ATP in a total volume of 50 µL. The capped and polyA-tailed transcripts were purified following the RNA Clean & Concentrator-5 protocol (Zymo Research) and eluted in 33 µL of nuclease-free water (NFW). From this sample, a volume of 3 µL was kept separately and used as a non-enriched control sample. In the enriched samples, the desthiobiotinylated primary transcripts were captured with Hydrophilic Streptavidin Magnetic Beads (New England Biolabs). For this, an equal volume of beads, prepared by washing three times in Washing Buffer (10mM Tris-HCl pH7.5; 250mM NaCl; 1mM EDTA) and resuspension in Binding Buffer (10mM Tris-HCl pH7.5; 2M NaCl; 1mM EDTA), was mixed with the RNA and incubated on a rotator at room temperature for 45 minutes. After washing three times in Washing Buffer, beads were suspended in 50 µL Biotin Buffer (1mM Biotin; 10mM Tris-HCl pH7.5; 50mM NaCl; 1mM EDTA) and incubated on a rotator at 37°C for 30 minutes to elute the RNA. The biotin was removed from the samples and the purified RNA was eluted in 10 µL NFW using the Zymo RNA Clean & Concentrator-5 kit. In parallel, the control samples were subjected to the same incubation steps, but the enrichment procedure was omitted.

#### Reverse transcription and selection for full-length transcripts

Reverse transcription and PCR amplification was performed according to the cDNA-PCR Barcoding protocol (SQK-PCS109 with SQK-PBK004; Oxford Nanopore Technologies), with an additional selection for full-length primary transcripts in the enriched samples. Briefly, 9 µL of enriched RNA (enriched sample) or 1 µL of non-enriched RNA (control) was mixed with oligo(dT) VN primers and 10 mM dNTPs in a total volume of 11 µL, incubated for 5 minutes at 65°C, and snap-cooled on ice. Next, first strand synthesis was carried out as described in the protocol with 200 units of Maxima H Minus Reverse Transcriptase (Thermo Fisher Scientific, followed by incubation for 90 minutes at 42°C and inactivation for 5 minutes at 85°C. The enriched samples were then treated with 50 units of RNase If (New England Biolabs) for 30 minutes at 37°C. All samples were purified using AMPure XP beads (Beckman Coulter) and subsequently eluted in Low TE buffer (1 mM Tris-HCl pH7.5; 0.1 mM EDTA). Before proceeding to second-strand synthesis, the enriched samples were subjected to a second round of streptavidin-based enrichment to select full-length cDNA molecules. For this, the cDNA/RNA duplexes were incubated again with an equal amount of prewashed Hydrophilic Streptavidin Magnetic Beads and incubated as stated before. The beads were washed three times with Washing Buffer and resuspended in 20 µL of low TE buffer. Afterwards, second-strand cDNA synthesis and PCR Barcoding was carried out on all samples using the manufacturers’ guidelines. For PCR barcoding, 16 cycles of amplification were performed, each with an extension time of 15 min to ensure full-length amplification.

#### Library preparation and nanopore sequencing on the PromethION platform

The cDNA reaction products were treated with 20 units of Exonuclease 1 (New England Biolabs) and subsequently purified and concentrated using AMPure XP beads. The quantity of the amplified cDNA barcoded samples was measured using the Qubit 1X HS dsDNA Assay kit (Thermo Fisher Scientific). Equimolar amounts were pooled to a total of 100 fmol, in a 23 µL sample volume. Finally, the cDNA library was mixed with nanopore sequencing adapters, loaded on a PromethION flow cell (R9.4.1) and run on a PromethION 24 device with live base-calling and demultiplexing enabled. After 48h, the flow cell was refuelled and reloaded with the amplified cDNA library, and the sequencing run was continued for an additional 48h until all pores were exhausted.

### 6. Data analysis

#### Raw read processing and QC

All fastq files that exceeded the default phred-like quality score threshold after base calling (>7) were used as an input for downstream data analysis. Overall sequencing performance, throughput and raw read quality was evaluated by NanoComp (v1.11.2) (27). Next, the raw reads were processed by Pychopper (v2.5.0) (https://github.com/nanoporetech/pychopper) to identify, rescue and correctly orient full-length cDNA reads. Cutadapt (v2.7) (32) was used to remove 3’ polyA stretches and sequence remnants on the 5’ end that match the primers used during reverse transcription, as described previously (16).

#### Read mapping to the reference genomes

The trimmed reads were mapped to the reference genomes of *Pseudomonas* phage LUZ7 (NC_013691.1) and *P. aeruginosa* strain US449 using minimap2 (v2.17) with settings recommended for nanopore cDNA reads (−ax -map-ont -k14) (33). To avoid spurious alignments, reads with more than 10 clipped bases at their 5’ and 3’ end were discarded from the SAM file using samclip (v0.4.0) (https://github.com/tseemann/samclip). SAMtools (v1.9) was used to convert the alignment format, generate sorted and indexed BAM files and evaluate the mapping metrics of each sample (34). Subsequently, the mapped reads were assigned to the genomic features of *P. aeruginosa* or LUZ7 using featureCounts (v2.0.1) in long-read mode (−L -O) (35). Alignments were visualized in the Integrative Genomics Viewer (IGV) software (36).

#### Identification of phage TSSs

The identification of transcriptional boundaries was carried out using custom scripts inspired by previously published work (16), with modifications tailored to our ONT-cappable-seq approach (https://github.com/LoGT-KULeuven/ONT-cappable-seq). For TSSs detection, the alignment files were used to generate strand-specific bed files that report the number of reads starting at each position in the LUZ7 genome using bedtools genomecov (−5 -bga) (37). Next, a peak calling algorithm (https://github.com/NICHD-BSPC/termseq-peaks) was applied to determine genomic positions of local maxima from 5’ read ends (16, 38). Nearby peaks within a distance of 5 bp on the same strand were clustered, retaining only the peak position with the highest number of 5’ read ends. Peak positions where less than five reads began were neglected.

In a second step, the read count per million mapped reads (RPM) was calculated for all remaining peak positions in the individual samples. To assess whether the accumulation of 5’ read ends arises from primary transcripts, an enrichment ratio was determined for each peak position i by dividing the RPM in the enriched sample by the RPM in the corresponding control sample with maximum 1 bp positional difference (i ± 1):

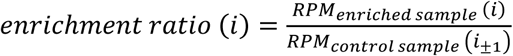

In case the enrichment ratio surpassed the threshold value, the peak position was annotated as a TSS. To accommodate for differences in enrichment levels across the samples, this threshold (T_TSS_) was set by assessing the enrichment ratio of the TSS of the T7 promoter of the RNA spike-in in the samples taken 5 minutes (T_TSS_ = 40.9), 10 minutes (T_TSS_ = 16.7) and 20 minutes (T_TSS_ = 24) post-infection. The -50 to +5 region of the annotated TSS was analysed using MEME (39) and late promoter regions (−100 to +1) were uploaded in SAPPHIRE (v2) (40) for *Pseudomonas* σ70 promoter sequence prediction.

#### Identification of phage TTSs

LUZ7 TTSs were pinpointed in a similar manner as the TSSs detection. In a first step, strand-specific bed files of the 3’ read ends were generated from the alignments of the enriched samples (bedtools genomecov -3 -bga) (37). Next, genomic positions with an accumulation of 3’ read ends were identified using the peak calling algorithm as described previously (16, 38). However, as ONT-cappable-seq does not allow direct differentiation between random termination events and true TTSs, more stringent filtering steps were implemented for 3’ end detection as opposed to TSS identification. Peak positions located on the same strand within 25 bp were clustered together. Here, the site with the largest number of reads was selected as a representative. Candidate TTSs with less than 20 reads ending at that position were filtered out.

In a second step, the strength of the terminators was estimated using custom scripts inspired by previously described methods (14, 41). Briefly, all reads that start upstream of the TTS were extracted and used to calculate the coverage drop before and after the terminator, averaged over a window of 20 bp. Only terminators that showed a read reduction of at least 20% were annotated as putative TTS and used for further analysis. Intrinsic transcriptional terminators were predicted using TransTermHP (42) and ARNold (43) after uploading the -50 to +50 region of the ONT-cappable-seq predicted TTSs. RNAfold (v2.4.13) was used to predict and calculate the secondary structure and minimum free energy of the -50 to +1 region of the TTS (44).

#### Determination of transcriptional units

The transcription units (TUs) were delineated based on adjacent TSSs and TTSs defined in this study. As a validation, we identified reads that encompass the candidate TUs in full-length. In case no clear TSS-TTS pair could be identified, the most distant 3’ end was used as the putative TU boundary. The unique combination of phage genes encoded in the transcription units, defined as the transcriptional context (15), was determined by finding overlaps between the ONT-cappable-seq reads and the genomic features of LUZ7 using the bedtools intersect tool (37). Only the genes encoded on the same strand that are least 90% covered by the transcription unit were included in the transcriptional context (−F 0.9 -s). Overlapping transcriptional units that were transcribed from the same strand and had at least one common gene in their transcriptional context were annotated in an operon structure (45).

### 7. Primer extension assays

Primer extension assays were carried using the Superscript IV First-Strand Synthesis System (Thermo Fisher Scientific), according to the manufacturer’s instructions. For this, specific primers (Supplementary Table S1) were radiolabelled with T4 Polynucleotide kinase (NEB) and [?-^32^P]ATP (6,000 Ci/mmol), purified on a Micro-Bio Spin P-6 Gel Column (Bio-Rad)and subsequently annealed to 5 µg of RNA extract. The RNA-primer samples were shortly incubated at 65°C, supplemented with the SuperScript IV Reverse Transcriptase reaction mix and incubated at 55°C for 10 minutes. The reaction product was mixed with formamide loading dye, boiled for 1 min and loaded on an 8% polyacrylamide-7M urea gel.

In parallel, for each TSS candidate, the surrounding DNA region was amplified from the LUZ7 genome using the primer for primer extension, paired with a primer located upstream the TSS of interest (Supplementary Table S1). The resulting PCR fragment was used as a template for Sanger sequencing to infer the nucleotide sequence of the transcription initiation region. The Sanger sequencing reaction was carried out using the USB Thermo Sequenase Cycle Sequencing Kit (Affymetrix), following the guidelines for radiolabelled primer cycle sequencing. Briefly, after mixing the radiolabelled primers, the DNA template, and a dNTP mix containing a dideoxy variant of one of the nucleotides (ddATP, ddTTP, ddCTP or ddGTP), four sequencing reactions were performed for 45 cycles (95°C, 30s; 55°C, 30s and 72°C, 1 min). The samples were supplemented with loading dye, boiled for 1 min and loaded on an 8% polyacrylamide-7M urea gel alongside their respective primer extension product. The gel was fixed, dried and exposed on a phosphor screen before visualising with a Typhoon phosphorimager (GE Healhcare).

### 8. In vivo promoter activity assay

The functionality of selected LUZ7 late promoters was confirmed experimentally *in vivo* using our recently established SEVAtile-based expression systems. For this, adaptors (generated by overlapping primer pairs) encoding putative promoters were cloned upstream of a standardized ribosomal binding site (BCD2) and an msfGFP reporter gene in a pBGDes vector using SEVAtile assembly (46). In addition, a vector lacking a promoter sequence (pBGDes BCD2-msfGFP) and a vector with a constitutive promoter (pBGDes P_EM7_-BCD2-msfGFP) was created to serve as a negative and positive control, respectively. As a separate negative control, we constructed three vectors with a random LUZ7 sequence as a decoy promoter site (pBGDes P_decoy_1-3-BCD2-msfGFP). Vectors, primers and insert sequences used are listed in Supplementary Table S1. The genetic constructs were transformed to both *E. coli* PIR2 and *P. aeruginosa* PAO1 host cells. In case of *P. aeruginosa* PAO1, the resulting vectors were electroporated together with a pTNS2 helper plasmid according to the method by Choi *et al* (2005) to ensure site-specific genomic integration (47). To quantify the *in vivo* expression levels of msfGFP in the transformed *P. aeruginosa* PAO1 and *E. coli* PIR2 cells, fluorescent expression assays were performed using four biological replicates. The bacteria were inoculated in M9 minimal medium (1 x M9 salts (BD Biosciences), 0.5% casein amino acids (LabM, Neogen), 2 mM MgSO_4_, 0.1 mM CaCl_2_ (Sigma Aldrich), 0.2% citrate (Sigma Aldrich)) supplemented with 50 µg/µL kanamycin (*E. coli* PIR2) or 30 µg/mL gentamicin (*P. aeruginosa* PAO1) and grown overnight. The next day, the cultures were diluted in fresh M9 medium with the appropriate antibiotic in a Corning® 96 Well Black Polystyrene Microplate with Clear Flat Bottom and incubated with shaking. After 3.5 hours, msfGFP and OD_600_ levels were measured on the CLARIOstar® *Plus* Microplate Reader (BMG Labtech, Ortenberg, Germany), as described previously (46). The msfGFP fluorescence units were normalized for the associated OD_600_ values and converted to absolute units using an independent calibrant, 5(6)- carboxyfluorescein (5(6)-FAM) (Sigma Aldrich) (46, 48).

### 9. In vivo terminator validation

A subset of the LUZ7 terminators was validated *in vivo* by a terminator trap system, as described by Lammens *et al*. (2021). Briefly, pBGDes vectors bearing the terminator regions of interest between an msfGFP reporter upstream and a mCherry reporter downstream were generated using SEVAtile assembly (46) (Supplementary Table S1). Expression of both reporters is driven by the P_EM7_ constitutive promoter and standardized translation initiation elements BCD1 and BCD2 (49, 50). Hosts carrying constructs without terminator (pBGDes P_EM7_-BCD2-msfGFP-BCD1-mCherry) and the wildtype T7 terminator from coliphage T7 (pBGDes P_EM7_-BCD2-msfGFP-T7(wt)-BCD1-mCherry) were used as a negative and positive control, respectively. Next, fluorescent expression assays were performed to quantify the transcriptional termination activity of the terminators of interest. Similar to the promoter activity assay described above, four biological replicates of *P. aeruginosa* PAO1 carrying the constructs were grown in M9 minimal medium with antibiotic and subsequently assayed for fluorescence measurements. After 3-4 hours incubation in fresh M9 medium the next day, the msfGFP, mCherry and OD_600_ levels were measured on the CLARIOstar device. The fluorescent intensity of msfGFP and mCherry was determined for each sample, and normalized for the corresponding OD_600_ value. The termination activity was subsequently calculated according to previously published methods (46, 51):

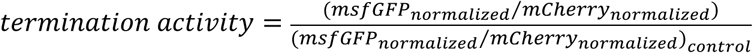

## RESULTS AND DISCUSSION

### 1. Profiling the full-length primary phage transcriptome using ONT-cappable-seq

In this study, we combine the Cappable-seq enrichment strategy and the nanopore sequencing platform to isolate and sequence the full-length primary transcriptome of *Pseudomonas* phage LUZ7 at different infection stages of its host. The key steps of the ONT-cappable-seq workflow are depicted in Figure 1a. The RNA transcripts are subjected to an enzymatic capping reaction that specifically labels the 5’ triphosphate group hallmarking the beginning of primary transcripts with a desthiobiotin tag (9). Next, the transcripts are polyadenylated to preserve their 3’ ends and make them compatible with nanopore RNA sequencing library preparation. To selectively enrich primary transcripts, the desthiobiotinylated and polyadenylated transcripts are captured on streptavidin beads and separated from their processed counterparts (9). Afterwards, the RNA is reverse transcribed and the second round of enrichment is performed to select for full-length transcripts (15). The cDNA is barcoded and PCR-amplified, followed by adapter addition to feed and translocate the cDNA fragments through the nanopore. In parallel, an unenriched control library is similarly prepared from the same RNA sample by excluding streptavidin enrichment steps from the workflow. Depending on the number of samples and desired read depth, the final pooled library can be sequenced on a MinION or a PromethION device.

**Figure 1.**
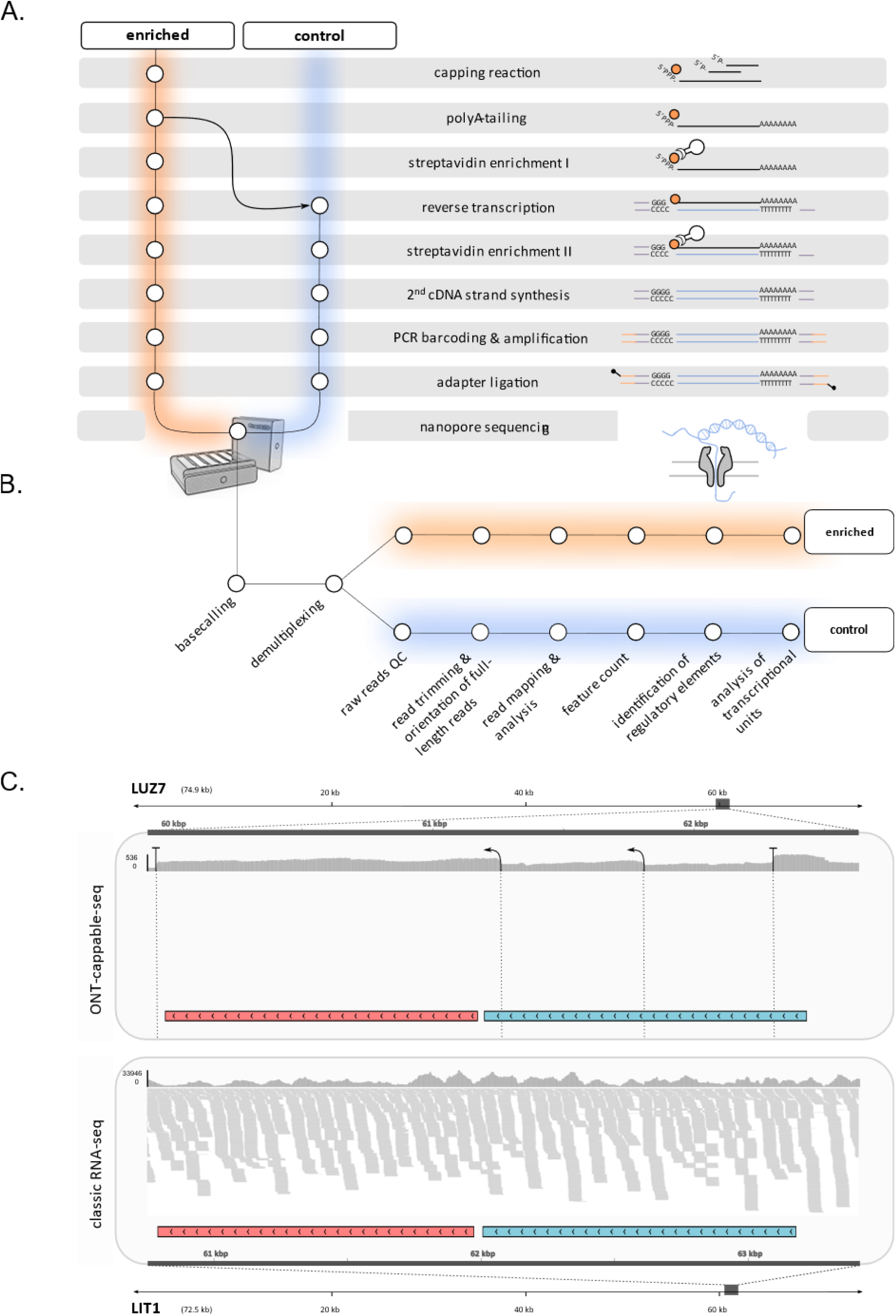
The ONT-cappable-seq technology. **a**. Schematic of the ONT-cappable-seq library preparation. After capping 5’ triphosporylated transcript ends with a desthiobiotin label, the primary transcripts are specifically enriched from the total RNA pool by streptavidin beads in the enriched library. The transcripts are polyA-tailed, converted to cDNA, barcoded and PCR amplified and loaded on the nanopore flow cell for sequencing. An unenriched control library is prepared in parallel. **b**. Bioinformatic pipeline implemented for LUZ7 phage transcripts analysis. The nanopore reads are base called and quality control is performed on the individual samples. Raw reads are trimmed, oriented and mapped to the reference genomes, followed by analysis of read distribution on genomic features and the elucidation of phage LUZ7 genome transcriptional architecture. **c**. Comparison of Integrative Genomics Viewer (IGV) data tracks between the late transcriptomes of LUZ7 and LIT1 that were sequenced using ONT-cappable seq (top) or Illumina-based RNA-seq technology (bottom), respectively. Only the region of phage genomes that comprise two highly conserved genes in N4-like phage members encoding the major capsid protein (red) and N4 gp57-like protein (light blue), are shown in the IGV representations. While ONT-cappable-seq can accurately define TSS (arrows) and TTS (T) in LUZ7, genuine transcriptional boundaries are more challenging to identify from LIT1 classic RNA-seq data.

To obtain temporal transcriptional landscapes of LUZ7 throughout infection of its *P. aeruginosa* US449 host, uninfected (0 min), early (5 min), middle (10 min) and late (20 min) infection stage samples were taken and, together with their corresponding controls, multiplexed and loaded on a single PromethION flow cell. The sequencing experiment ran for >48 hours, generating a total of 43.3 million reads (18.2 Gb) that passed the default quality score threshold after base calling (Supplementary Figure S2). The full-length reads were trimmed, oriented and mapped against the reference genomes of the phage and the host for further downstream analyses (Figure 1b). The number of reads that could be mapped to both genomes ranged between 69.9% and 78.4% in the individual samples (Supplementary Figure S3a).

The enrichment of primary transcripts is a critical step in the ONT-cappable-seq method and can be evaluated by analysing the fraction of reads that originate from processed RNA species, such as ribosomal RNA (rRNA) and transfer RNA (tRNA) (15, 52). We compared the distribution of reads on the genomic features of LUZ7 and *P. aeruginosa* US449 between the enriched and the control libraries of each dataset. We observed that more than 97.5% of the mapped reads in the control libraries correspond to rRNA and tRNA. In contrast, the number of reads from rRNA and tRNA was significantly reduced (28.6%-60.4%) in the enriched libraries of each timepoint, indicating successful selection against processed transcripts (Figure 2). After sequencing the enriched samples, 96-99% of the annotated genes are covered by at least one read in all datasets, whereas only 65-80% of the genes are covered in the control samples. This shows that the enrichment procedure and the reduced number of processed RNA species enables deeper sequencing of the prokaryotic transcriptome.

**Figure 2.**
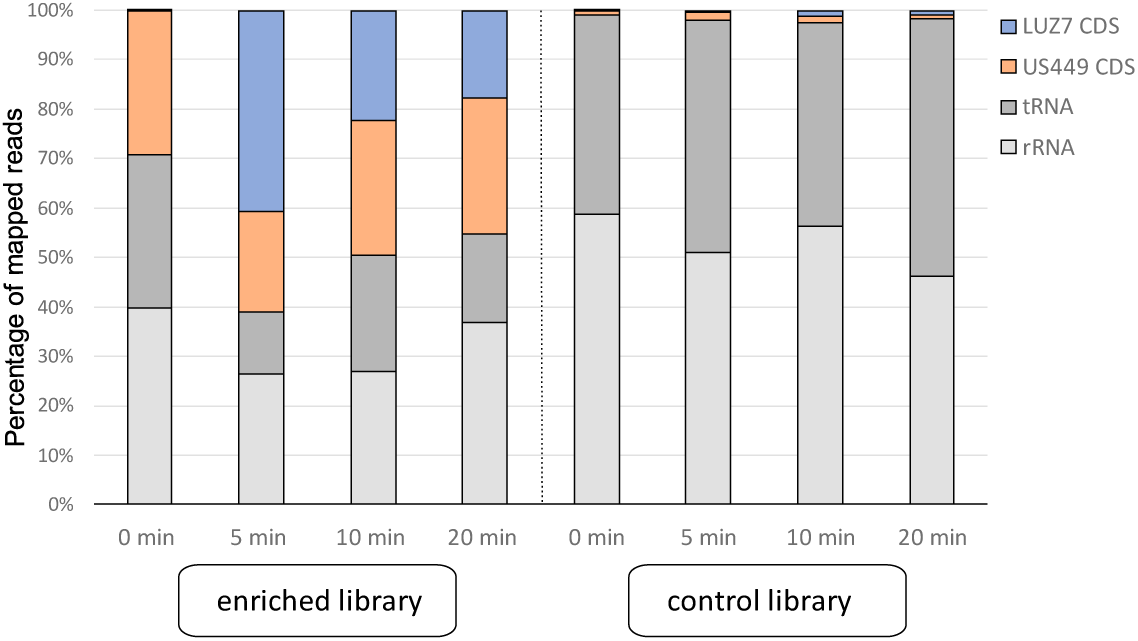
Distribution of reads across the genomic features of phage LUZ7 and its *P. aeruginosa* US449 host. The percentage of reads mapped to rRNA (light grey), tRNA (dark grey) and the coding sequences (CDS) of *P. aeruginosa* US449 (orange) and phage LUZ7 (blue) for each timepoint for enriched and control libraries is presented.

To further evaluate whether the ONT-cappable-seq procedure provides a comprehensive picture of time-resolved full-length transcriptional landscape during phage infection, we analysed the length of the mapped reads in individual samples (Supplementary Figure S3b). As expected, the aligned read length distribution was impacted significantly by the enrichment procedure with N50 read lengths varying between 358-372 bp for the enriched samples and 1,511-1,541 bp for the controls. The length difference is more pronounced on the bacterial side and can be largely attributed to rRNA depletion, as the mean reads lengths and N50 values of phage-derived transcripts are similar between enriched and control libraries across different samples. Despite modest average read lengths, the top five longest mapped phage reads in each sample span 3.3 kb-8.3 kb, indicating that the procedure allows end-to-end sequencing of long RNA molecules. The advantage of this long-read RNA sequencing method to recover the boundaries of individual phage transcripts is illustrated when comparing the Integrative Genomics Viewer (IGV) data tracks of our technique to standard RNA-seq experiments performed previously on another N4-like phage, LIT1 (24) (Figure 1c).

### 2. Delineation of the transcriptional features of *Pseudomonas* phage LUZ7

ONT-cappable-seq enables full-length profiling of primary transcripts, providing information on their 5’ and 3’ extremities simultaneously. sing this approach, we generated a comprehensive inventory of phage LUZ7 transcripts, unveiling transcription initiation and termination events, complex operon structures and gene expression levels across different infection stages, all in a single experiment (Figure 3).

**Figure 3.**
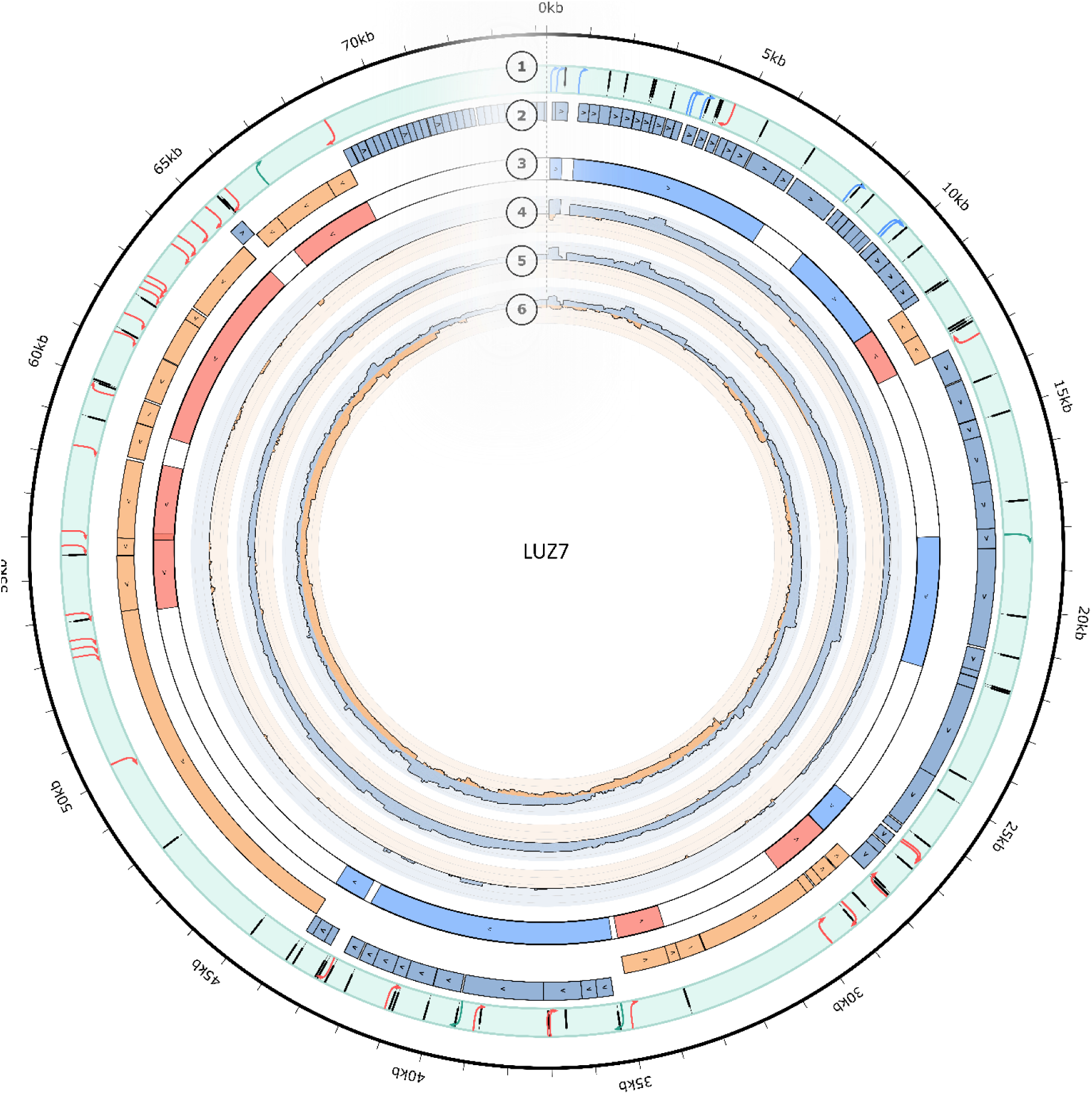
Circular representation of the genome and transcriptional landscape of phage LUZ7. The outer circle displays the position and orientation of transcriptional initiation sites (arrows) and termination sites (bars) identified by ONT-cappable-seq. The TSS are marked in blue (early promoters), green (middle promoters), or red (late promoters). Circle 2 depicts the genome annotation of LUZ7, showing CDS annotated on the Watson strand in blue, and CDS encoded on the Crick strand in orange. It is followed by a stranded representation of the complex operon structures identified in this study (circle 3). The orientation of the operons is indicated by colour (blue = Watson strand, red = Crick strand) and an arrow showing the direction of transcription. The tree inner circles represent the normalized log10 expression profiles 5 (circle 4), 10 (circle 5), and 20 minutes (circle 6) post-infection. Genome-wide read coverage of the Watson and Crick strand is indicated in blue or orange, respectively. The figure made created using the Circos visualization tool on the Galaxy platform (53).

#### Identification of LUZ7 transcription start sites

Candidate transcription start sites (TSSs) are identified by determining the genomic positions where a significant number of reads begin. Afterwards, comparison of the relative strength of each TSS candidate between the enriched library and the control reveals the positions of original 5’ transcript ends, enabling TSS determination at high resolution. TSS identification by our method was first verified using an *in vitro* transcribed RNA spike-in, which was added to RNA from phage-infected cells prior to library preparation (Supplementary Figure S4). This demonstrated accurate TSS annotation of the spike-in transcript initiated from a T7 RNA polymerase promoter. Next, the boundaries of LUZ7 own transcripts were delineated. Comparative transcriptomic analysis, followed by manual curation, yielded a total of 46 unique TSSs across the LUZ7 genome, which were classified according to their genomic context and expression level as described previously (8) (Supplementary Table S2). Primary TSSs are defined as TSSs that reside within ≤500 bp upstream of annotated genes and show a higher expression level than additional TSSs for the same gene (secondary TSSs). Intragenic TSS on the same strand of an encoded gene, antisense TSS located inside or within 100 bp distance of an oppositely oriented gene, and TSS with no genes nearby are designated as internal, antisense, and orphan TSS, respectively. In this manner, we observed 25 primary and three secondary phage TSS, and found a considerable overlap between the five different TSS categories (Figure 4a).

**Figure 4.**
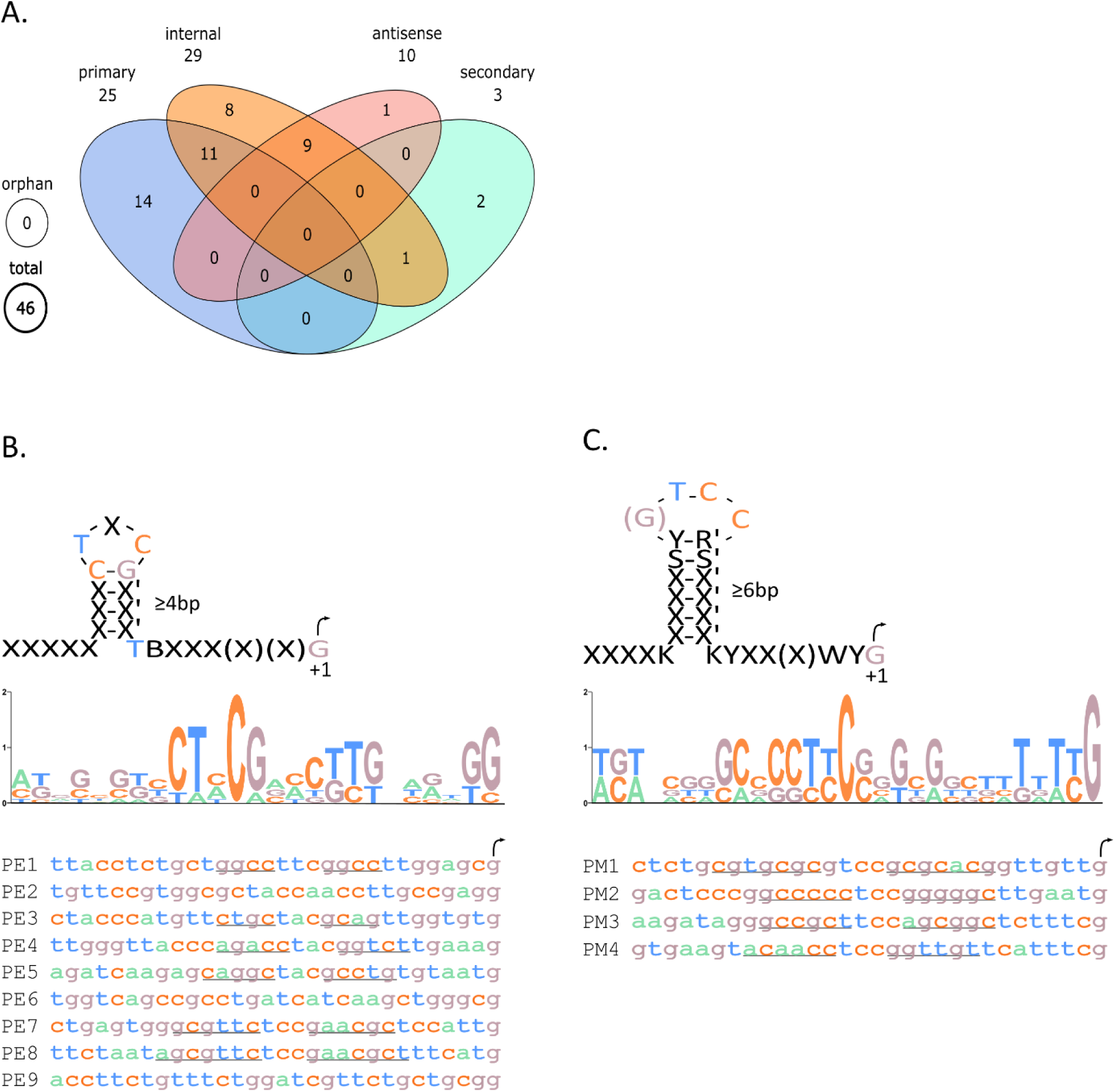
TSS identification in LUZ7. **A**. Venn diagram showing viral TSS classifications. **b**. Motif analysis of the early promoters reveals a conserved hairpin structure similar to that found in N4. **C**. Motif analysis of LUZ7 middle promoters reveals a GC-rich hairpin structure.

Multiple sampling points throughout the LUZ7 infection allowed us to distinguish between early, middle, and late phage promoters, from which three different RNAPs successively initiate transcription during infection. Transcription of the LUZ7 genome begins after the phage co-injects its viral DNA along with a large vRNAP that drives transcription from early phage promoters. After analysing the viral primary transcriptome during the early infection stage, nine TSSs and their cognate early promoter sequences were identified, including two promoters that were previously predicted after inspection of the LUZ7 genome (21). The majority of early promoters contain a typical hairpin structure similar to that of coliphage N4 early promoters (Figure 4b), with a ≥4 base pairs (bp) stem and a conserved three nucleotide loop (54). Motif analysis revealed that the preferred TSS nucleotide of the identified early promoters is guanosine, located 13-14 bp downstream of the hairpin centre. Notably, the few early promoters that lack the distinctive hairpin structure all reside within the coding region of genes that are preceded by a hairpin promoter, yielding truncated transcripts. These non-canonical early promoters may therefore depend on a prior initiation from their canonical hairpin counterpart, though this requires further investigation.

Interestingly, two of the strongest early TSSs are organized in tandem, 33 nucleotides apart, and drive the expression of early phage protein Drc, making it the most highly expressed viral gene during infection, accounting for up to 64% of phage transcripts. The Drc protein was recently shown to be a key transcriptional regulator during LUZ7 infection, activating the progression from the early to the middle transcription stage by recruiting the LUZ7 RNAPII to middle promoters (22). In addition, *in vitro* this protein can fully cover large stretches of (ss)DNA when present at high concentrations and since its transcripts are so abundant this is likely also the case *in vivo* during phage infection.

Middle promoters have thus far been impossible to identify in LUZ7 and related phages such as LIT1, despite the availability of middle stage transcription initiation sites in the prototypical phage N4 and RNA-seq data on phage LIT1 (24, 55). Our ONT-cappable-seq data on RNA from the mid-infection stage did enable us to identify four viral TSSs associated with middle stage infection. Collectively, these middle promoters show a GC-rich hairpin structure that resembles that of the early hairpin promoters, with a stem of ≥6 bp and a 3-4 bp nucleotide loop positioned ∼15 bp upstream of the identified TSS (Figure 4c). Similarly to the early TSSs, guanosine is the preferred nucleotide for middle promoters. The architecture of middle promoters is significantly different from the RNAPII transcription initiation sites that have been documented for coliphage N4, suggesting that LUZ7 and N4 have distinct transcriptional initiation mechanisms on their middle promoters, as in fact was proposed previously (22). In N4, RNAPII prefers to start transcription from thymidine and recognizes promoters with two consensus motifs in a sequence-specific manner, whereas transcription initiation by 7’s NAPII appears to be directed by the recognition of hairpin structures (20). These results indicate that ONT-cappable-seq can help infer the requirements of RNAPII promoter recognition in N4-like phages, which has proven to be very challenging in the past. However, given how extensive the middle genomic region is, it can not be excluded that the limited set of identified LUZ7 middle promoters is an underestimation of the true number of TSS associated with the middle transcription stage.

The third and final transcription stage of LUZ7 is driven by the host RNAP, which is recruited to late promoters on the phage genome. We discovered 33 late TSSs scattered across the LUZ7 genome, of which more than 80% are located on the Crick strand. This strand bias is largely in agreement with the orientation of the structural and lysis gene cassettes, which is opposite to that of the early and middle genes. Notably, all genes on the Crick strand are strictly transcribed during the late stage, even when these genes are found inserted within early gene cassettes (e.g. genes *orf34* and *orf35*). In N4, late promoters show remote resemblance to the σ^70^ promoter sequences of the host. To assess whether this is also true for LUZ7, nucleotides -100 to +1 bp of the identified late TSSs were analysed using the *Pseudomonas* promotor prediction tool, SAPPHIRE.CNN (40). Indeed, the vast majority of late promoters displayed significant similarity to the *Pseudomonas* σ^70^ promoter consensus sequence.

##### Experimental validation of viral TSSs and associated promoter sequences

To further benchmark the ability of the ONT-cappable-seq method to identify TSSs and promoter sites, a selection of the TSSs was validated individually. RNA extracted from early and middle infection stages was used to map representatives of the early (PE3) and middle (PM2) promoters and their associated TSSs using primer extension assays, and a near exact match with data obtained from ONT-cappable-seq analysis was obtained (Figure 5a).

**Figure 5.**
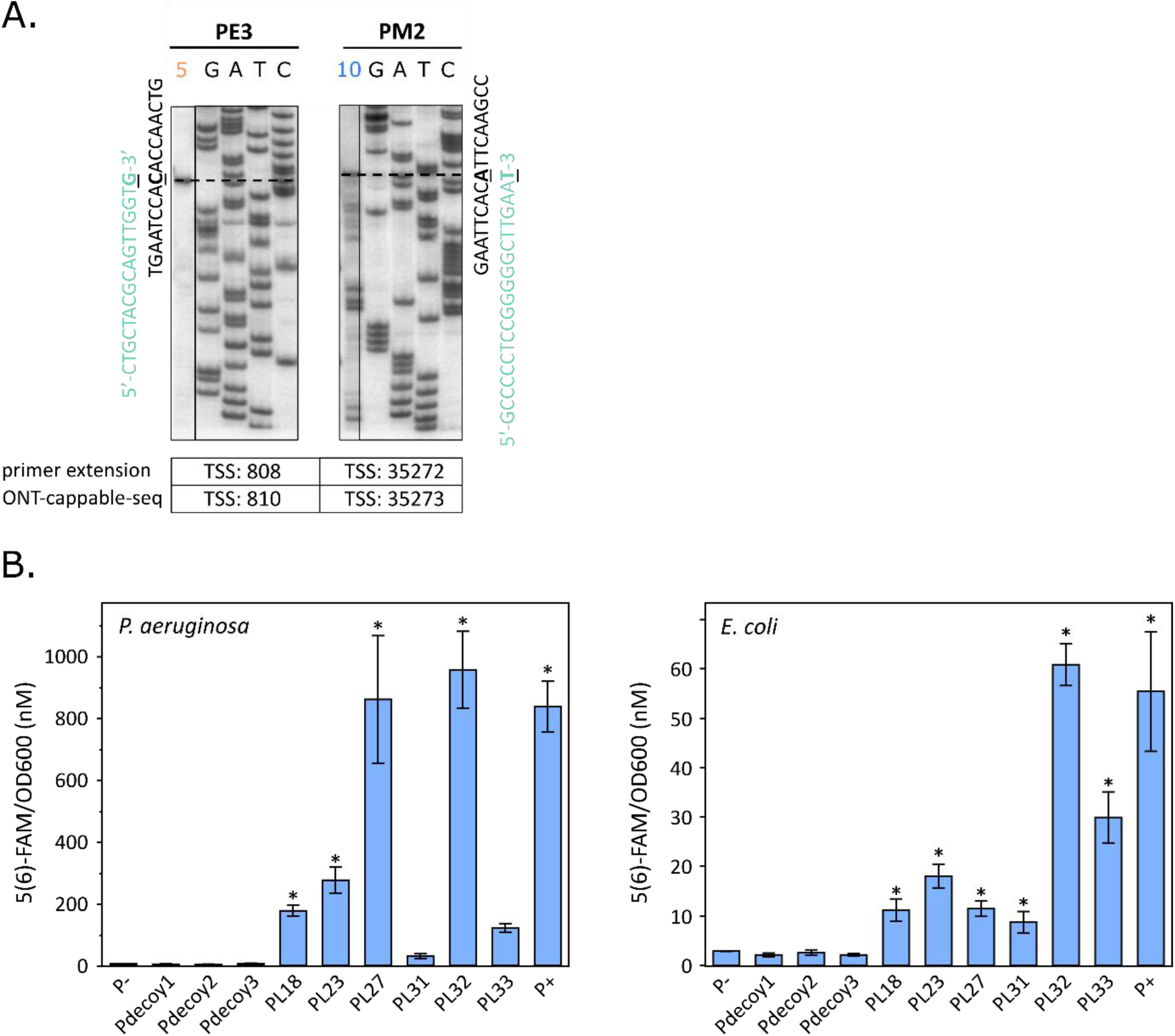
Experimental validation of TSSs and promoters identified by ONT-cappable-seq. **a**. Primer extension assays of early (PE3) and middle (PM2) promoter representatives. The products of primer extension reactions performed with RNA prepared from cells collected 5 (early stage of infection) and 10 (middle stage of infection) minutes post-infection were resolved by denaturing polyacrylamide electrophoresis next to appropriate Sanger sequencing reactions markers. The dashed lines represent the mobility of the primer extension end product band. The corresponding sequence is depicted next to the gel with the mapped TSS indicated in bold and underlined. Due to the orientation of the primer, this sequence represents the reverse complement of the coding strand sequence, which is provided in green alongside it. The table depicted below indicates the genomic coordinates of the TSSs as identified by primer extension and ONT-cappable-seq. **b**. *In vivo* validation assays of promoter activity in *P. aeruginosa* (left) and *E. coli* (right) of a subset of LUZ7 late promoters using SEVAtile-based expression systems. The negative control (−) represents a vector without promoter sequence and in the positive control (+) the msfGFP reporter is transcribed from a constitutive promoter (Pem7). Pdecoy1-3 are constructs containing a random LUZ7 sequence as promoter insert. The promoter activity was quantified by measuring the fluorescence intensity level normalized based on OD600 values and converted to absolute units of 5(6)-FAM fluorescence (displayed in 5(6)-FAM/OD600 nM). Promoters showing significantly higher fluorescence intensity compared to the negative control are indicated with an asterisk (*) (Student’s t-test, p < 0.05). Data represent the mean value for five replicates and standard deviation is indicated by error bars.

As a separate validation, the activity of a subset of the late LUZ7 promoters was evaluated *in vivo* using a promoter trap system using the SEVAtile-based expression systems (46). For this, genetic constructs containing a late phage promoter, a standard ribosome binding site (RBS), and the green fluorescent reporter protein (msfGFP) gene were constructed with the SEVAtile assembly method, followed by transformation to *P. aeruginosa* and *E. coli*. If the identified phage promoters are recognized by the σ^70^ factor of the host RNAP holoenzyme as predicted, expression of the fluorescent reporter gene shall be detected. Indeed, in *E. coli*, all promoters show elevated levels of msfGFP fluorescence intensity compared to constructs with a random LUZ7 sequence (P_decoy_1-3) and the negative control (P-), which lacks a promoter sequence insert (Student’s t-test, p < 0.05) (Figure 5b). Interestingly, while the majority of promoters seem to behave relatively similar in *E. coli* and *P. aeruginosa*, PL31 and PL33 exhibited a decreased activity in *P. aeruginosa* and do not promote significant expression of msfGFP in this genetic context. In addition, PL32 appears to have the strongest *in vivo* activity in both hosts, whereas PL27 shows relatively stronger expression levels in *P. aeruginosa* compared to *E. coli*. Be that as it may, together with the primer extension validations, these findings highlight the potential of our approach to carefully pinpoint phage transcription initiation events in a systematic manner across the genome.

#### Identification of phage transcription termination sites

In addition to TSSs, the annotation of 3’ transcript ends is essential to obtain a complete picture of an organisms’ transcription. The longer reads from nanopore-based sequencing allow the detection of the 3’ ends. However, the determination of true TTSs is often blurred by the activity of 3’ to 5’ exonucleases that obfuscate the true 3’ boundaries of NA molecules. As some of the major bacterial ribonucleases show a substrate preference for transcripts with 5’ monophosphate termini, primary transcripts with intact 5’ triphosphate groups are less susceptible to NA processing and are more likely to contain their original termination ends (56, 57). Consequently, the enrichment of primary transcripts performed in this study increases the likelihood that the observed termination signals represent genuine TTS, but it cannot be excluded that a number of 3’ ends originate from post-transcriptional processing.

In bacteria and phages, the termination of transcription largely occurs via two main mechanisms, intrinsic termination and factor-dependent termination. In general, transcription termination motifs were located on the phage genome by detecting local maxima of 3’ read ends in our enriched datasets, revealing a total of 61 termination sites, among which 14 are predicted to be intrinsic, factor-independent terminators (Supplementary Table S3). The vast majority of identified TTSs (65.6%) end transcription of RNA molecules transcribed from the Watson strand, whereas 21 of the termination signals are encoded on the Crick strand. Almost half of the termination sites (47.5%) are located in intragenic regions, directly downstream of the preceding gene on the same strand. The rest of the phage TTSs reside either in intergenic regions (32.8%) or in antisense orientation relative to annotated genes (19.7). The length of the 3’ untranslated regions (3’ UTR) varies extensively, with more than 30% of the 3’ UTR s exceeding 100 nt in length (Supplementary figure S5a). Analysis of regions upstream (−50 to +1) of TTS indicated that most phage terminators are prone to form stable secondary structures (Figure 6a). For each terminator, the termination efficiency (TE) was determined by calculating the level of readthrough across the termination site, and a wide range of terminator strengths was observed (Supplementary figure S5b). On average, the termination efficiency is weakly correlated with the folding energy of RNA upstream of the TTS sequence (R = -0.252, p < 0.05) (Supplementary Figure S5c). Interestingly, ten terminators showed altered readthrough levels (≥30%) throughout the infection. This suggests that these dynamic transcriptional terminators may help tweak LUZ7 gene expression levels throughout the infection (Figure 6b,c). For example, the TE of terminator T8 is less than 5% at the early infection stage, allowing almost all transcripts to fully span the early transcribed *orf15*, whereas 10 minutes post-infection the TE of T8 increases and almost 50% of the transcripts end prematurely within the *orf15* gene body. Similarly, the TE of terminator T42 decreases from 88% in the middle stage of infection to 57% at the late infection stage, enabling more transcripts to read through to gene *orf65*, which codes for an important regulator in the final transcriptional stage of LUZ7. Modulation of transcription termination is a powerful strategy to regulate gene expression programs and has also been investigated in detail in a handful of phages, including coliphages λ and HK022 and *Thermus thermophilus* phage P23-45 (58–61). The phage-encoded antiterminators manipulate the host RNAP and enable it to bypass terminators at the appropriate transcription stage during infection. In LUZ7, the varying terminator read-through levels throughout infection might point to similar or novel antitermination strategies that successively modifies the different RNAPs during phage transcription in response to regulatory stimuli, though this warrants further investigation.

**Figure 6.**
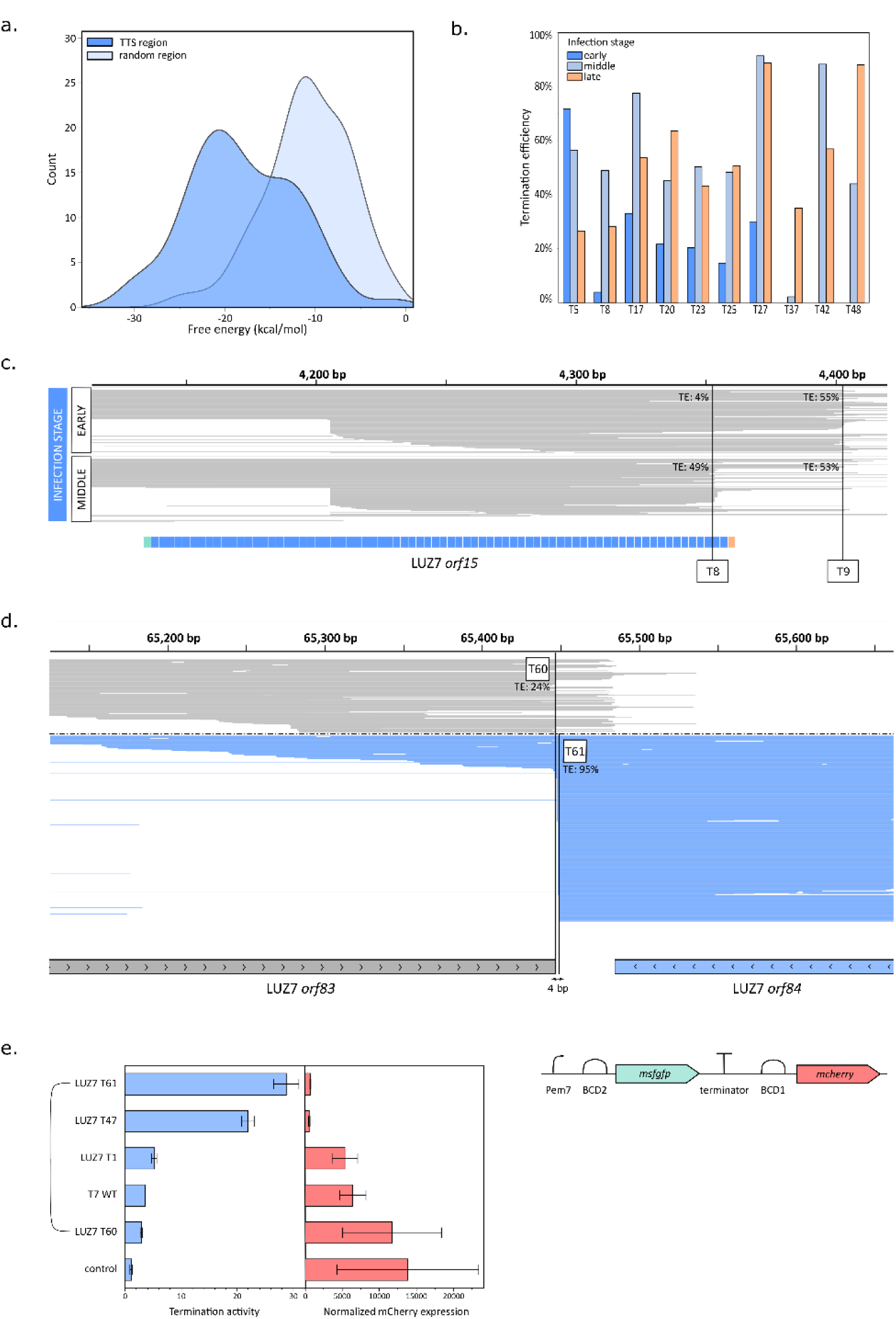
Characteristics of LUZ7 TTS and terminators. **a**. Distribution of the minimum free energy (kcal/mol) of the -50 to +1 sequences of identified phage TTS (dark blue) compared to sequences of the same length randomly selected across the LUZ7 genome (light blue). **b**. Bar plot showing the termination efficiency of a set of phage terminators detected at the early (dark blue), middle (light blue) and late (orange) infection stages. **c**. Comparison of ONT-cappable-seq data tracks of early and middle infection for phage terminators downstream of LUZ7 *orf15*. The fraction of transcripts that span the entire length of the gene is significantly decreased at middle infection stage. **d**. ONT-cappable-seq data track of the late infection stage showing an example of a bidirectional terminator region located between a convergent gene pair. **e**. *In vivo* validation of LUZ7 terminators in *P. aeruginosa* using a terminator trap assay (46, 51). The terminator of interest is trapped between an msfGFP and a mCherry reporter, which are transcribed from a constitutive promoter (Pem7). BCD1 and BCD2 are used as ribosomal binding sites to initiate translation of the reporters. The control represents a terminator trap vector without a terminator sequence. The termination activity was quantified by comparing the ratio of the normalized msfGFP and mCherry expression levels with the ratio normalized msfGFP and mCherry levels in the control (Methods). The bidirectional LUZ7 terminator pair is connected with a bracket. Data represent mean values for four replicates and standard deviation is indicated by error bars.

In addition to unidirectional transcriptional termination events, we discovered four overlapping bidirectional terminator regions comprised of pairs of convergent TTSs located within 4-40 nt from each other (Figure 6d). All but one of these bidirectional terminators reside between genes that are oriented in a head-to-head manner. Comparison of the read-through levels in both directions suggests that termination tends to be more efficient in the orientation of the gene that is predominantly expressed (Supplementary figure S5d).

##### Experimental validation of LUZ7 terminator regions

The potential of ONT-cappable-seq to uncover TTSs and terminators was supported by validating the termination activity of a subset of the LUZ7 terminators, including T1, T47 and bidirectional terminator pair T60 and T61 in *P. aeruginosa*. To this end, we constructed a terminator trap using the SEVAtile assembly method, in which the phage terminators are inserted between two fluorescent reporter genes, *msfGFP* and *mCherry*, whose expression is driven by a constitutive promoter (46). To quantify termination activity, the intensity of msfGFP and mCherry expression was measured for all samples, and compared to control constructs containing either no terminator sequence or the native T7 terminator of phage T7 between the two genes, as described earlier (46, 51). Relative to the control, all phage terminators show reduced levels of downstream mCherry expression and, therefore, significant termination activity (Student’s t-test, p < 0.05), confirming their functionality *in vivo* (Figure 6e). These results further support the assumption that most TTSs annotated in this work represent genuine terminators.

#### Analysis of the operon organization

Taking advantage of the potential of ONT-cappable-seq to uncover both ends of transcripts, the operon architecture of LUZ7 can also be elucidated. In general, the boundaries of transcription units (TUs) were annotated based on adjacent TSSs and TTSs defined in this study. Next, the candidate TUs were validated by identifying ONT-cappable-seq reads spanning the full-length of a TU, leading to a total of 86 unique TUs (Supplementary Table S4). The length of the TUs varied extensively and it was found that LUZ7 TUs cover, on average, 1.9 genes (Supplementary Figure S6a,b). Analysis of the transcriptional context of TUs, defined by the unique combination of encoded genes (15), revealed significant overlap between individual TUs, as more than 60% of the genes in the annotated TUs are transcribed in more than one context (Figure 7a). This suggests that the alternative usage of TUs could serve as an additional regulatory layer to finetune the expression of individual genes (62).

**Figure 7.**
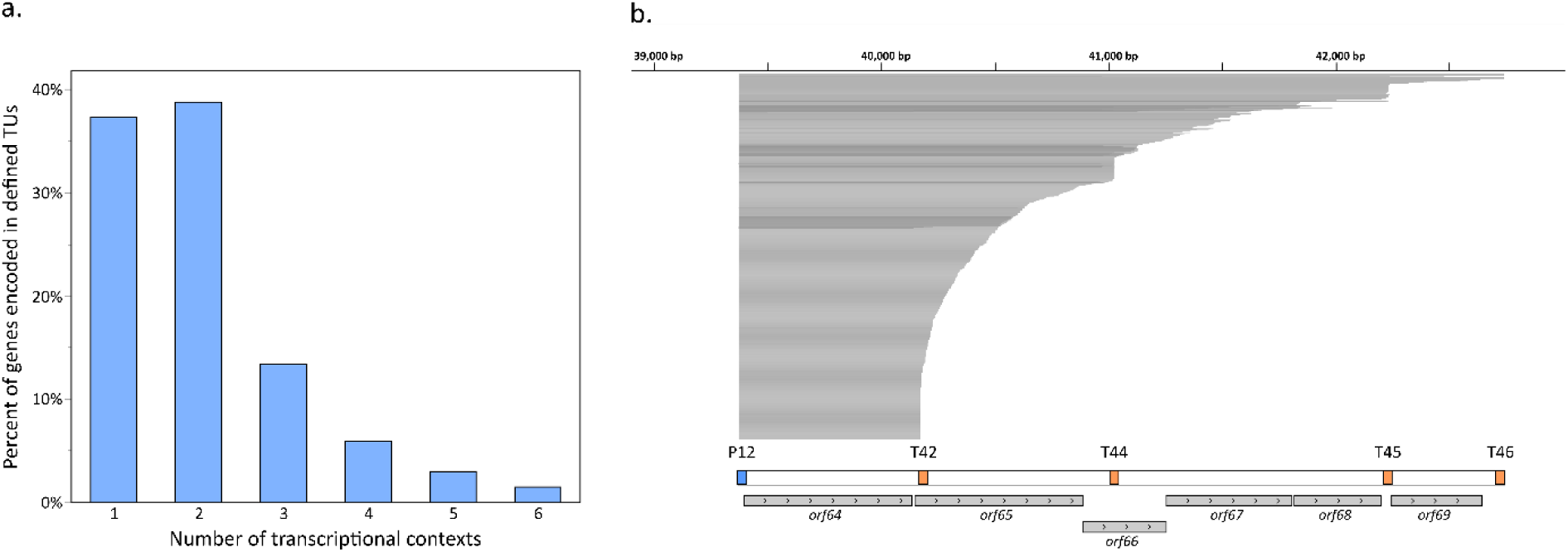
Transcriptional units of LUZ7. **a**. A bar plot showing the number of transcriptional contexts for genes encoded in identified transcriptional units. For each annotated gene, the number of transcriptional contexts represents the number of unique gene combinations within TUs that include that gene. Only genes that are transcribed in a least one transcriptional context are included. **b**. An ONT-cappable-seq data track showing the transcriptional pattern at LUZ7 genes *orf64*-*orf69* as an example of sequential readthrough at different termination sites for reads starting at the same TSS and leading to transcripts with up to five additional genes.

In the majority of cases, we observed that the gene content of transcripts altered as a result of sequential read-through across different TTSs, as exemplified by the stepwise transcription termination pattern of LUZ7 genes coding for gp64-gp69 (Figure 7b). This is consistent with the finding that more than 60% of the identified phage termination sites allow substantial downstream transcript extension (average TE <50%). Previous studies have already showcased that modulation of the degree of read-through at termination sites gives rise to numerous complex operon structures in bacteria (15). In addition, the use of inefficient terminators as a transcriptional strategy to properly balance gene expression levels has also been demonstrated for model coliphage T7 (63).

Based on the proposed definition of a complex operon as a gene cluster of which the transcription is orchestrated by various overlapping TUs that have at least one gene located on the same strand in common (45), the LUZ7 genome can be roughly organized in 14 complex operons, which are illustrated in summarizing Figure 3 (Supplementary Table S5). In general, genes that are included in the same operon structure are likely to be functionally related. This observation also appears to apply for LUZ7. For example, the first three operons residing in the leftmost genomic region comprise most of the early phage ORFans involved in the host hijacking process, whereas operons 7-8 and operon 13 encompass structural genes involved in the formation of the tail and the virion capsid, respectively (21, 24). It should be noted that not all annotated genes are included in identified operons. For example, a gene of the tail fiber protein (*orf*56, 3.2 kb) and of the virion-associated RNAP gene (*orf*073, 10.2 kb), two of the largest genes encoded by the phage, are not found in detected operons. This could indicate that unusually long transcripts pose more of a challenge to be fully captured by ONT-cappable-seq, making it more difficult to detect transcriptional unit variants that extend beyond these gene borders.

#### Identification of other regulatory features

In addition to cataloguing regulatory elements of LUZ7, ONT-cappable-seq allowed us to reveal peculiar transcriptional patterns of the phage. At the initial stages of infection, the genes encoding for hypothetical protein gp01 and the Drc protein (gp14) are intensively transcribed, reaching a combined total of 90% of early phage transcripts. Both gp01 and Drc have homologues in a number of N4-like *Pseudomonas* phages, including phage LIT1 (22, 24). Former analysis of the LIT1 transcriptome five minutes after *P. aeruginosa* PAO1 infection found comparable expression levels for the corresponding genes (24), corroborating that the encoded proteins play an important and seemingly conserved role in the infection strategy of N4-like phages targeting *Pseudomonas*. Whereas the crucial involvement of Drc in the transcriptional regulation of LUZ7 has already been demonstrated (22), a function for gp01 has not yet been identified.

Consistent with the transcriptional landscape of LIT1, we observe that the LUZ7 genome is almost exclusively transcribed from the Watson strand during early and middle timepoints (24). Interestingly, the transcripts captured mid-infection collectively cover the entire phage genome, including the rightmost region that encodes 29 small ORFans (gp87-gp115) (21). While the function of these genes is unknown, the temporal transcriptional information gained in this study could provide clues on the role of this unique gene cluster during the phage cycle. Given that the middle genes of LUZ7 are primarily involved in phage genome replication, these genes are likely to have a related function. In addition, ONT-cappable-seq data show that middle transcripts extend throughout genomic regions that have late structural gene cassettes encoded on the opposite strand. The same phenomenon was observed in LIT1, where it was suggested that the antisense transcripts are responsible for silencing premature expression of the late genes (24). During the final infection stages, the dominant transcription direction switches in both phages. Presumably, the surge in RNA molecules transcribed from the Crick strand overpowers the antisense transcripts to enable the timely expression of structural and lysis-associated genes (24).

Our data also revealed interesting transcriptional activity in the intergenic space between LUZ7 genes *orf69* and *orf70*. This region is extensively transcribed during the late infection stage, giving rise to numerous RNA species of varying lengths (Figure 8). Most of the reads that align with this region are less than 250 nt long and lack any apparent protein-coding potential, suggesting the presence of putative small non-coding RNAs (sRNA) in the LUZ7 transcriptome. Closer examination of this condensed transcriptional pattern roughly distinguishes three abundant sRNA candidates, designated as sRNA1, sRNA2, and sRNA3, that partially overlap with each other. Two of these RNA species share the same 5’ extremity but use an alternative termination site, resulting in a short and extended version of the transcript. By contrast, the shortest sRNA candidate, sRNA3, seemingly lacks an annotated TSS, suggesting that this fragment is derived from a primary precursor through processing events. Functional sRNA biogenesis through RNase activity has already been demonstrated in various bacteria (64–66), and was recently also suggested to occur in *Pseudomonas* phage phiKZ (67). In addition, the biological relevance of small non-coding RNAs in the regulation of phage and host development is increasingly being recognized in several other *Pseudomonas* phages (5, 68). However, in LUZ7, the regulatory scope of the putative sRNA species and the exact mechanisms that underlie their biogenesis remain to be elucidated.

**Figure 8.**
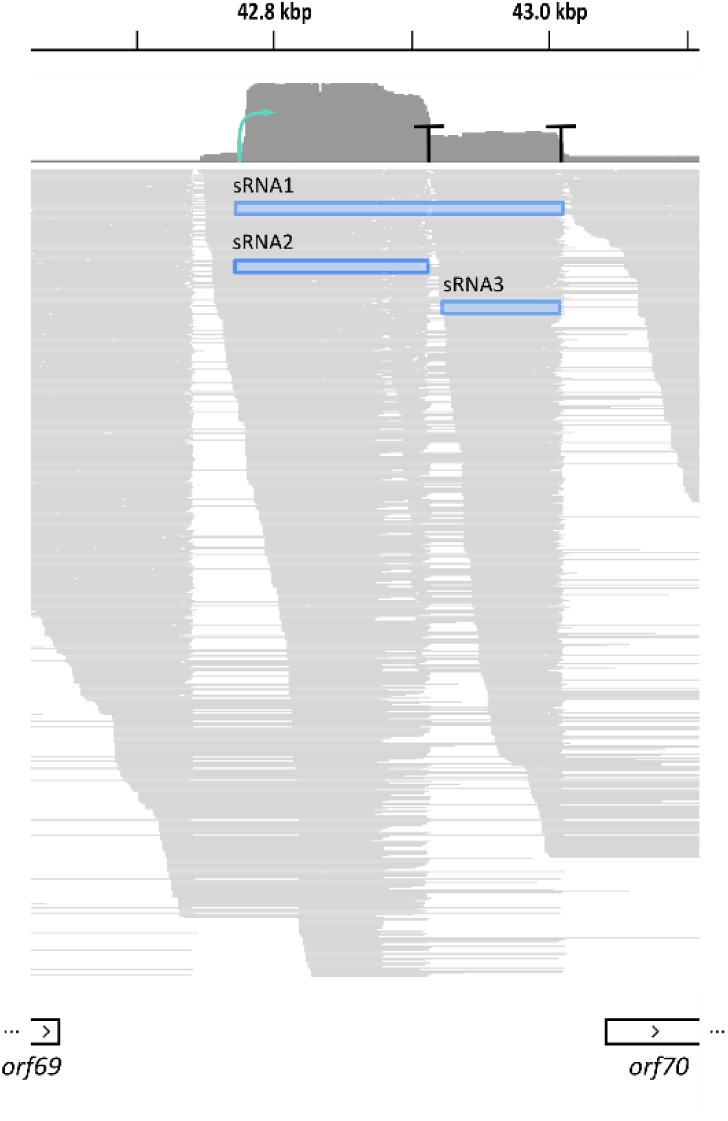
Unusual intergenic transcription activity during the late infection stage of LUZ7 reveals putative small non-coding RNAs. Visualization in IGV of late infection transcripts mapping to the intergenic space between LUZ7 genes *orf69* and *orf70* reveals a condensed cluster of small transcripts, suggesting the presence of putative small non-coding RNAs. The most abundant RNA species are highlighted with blue bars and denominated as sRNA1, sRNA2, and sRNA3. sRNA candidates sRNA1 and sRNA2 share the same TSS (indicated with a green arrow) but use a different terminator sequence (indicated with a ‘T’). The sRNA3 lacks an annotated TSS and may arise due to processing events.

## CONCLUSIONS AND PERSPECTIVES

The plethora of high-throughput microbial transcriptomic approaches that have emerged in the past decades has led to major advances in our understanding of the different layers of gene expression in bacteria. However, in contrast to bacterial RNA biology, the diverse transcription strategies of phages still remain poorly understood. We here took advantage of the Cappable-seq enrichment strategy and the nanopore sequencing platform to develop ONT-cappable-seq and generated a comprehensive transcriptional map of an N4-like *Pseudomonas* phage LUZ7. We showed that ONT-cappable-seq allows full-length transcriptional profiling and enables the simultaneous identification of both 3’ and 5’ transcriptional boundaries of individual phage transcripts. Using this approach, we pinpointed key phage-encoded transcriptional features at the early, middle, and late infection stages, unveiled complex operon organizations and revealed the presence of novel potential regulatory elements, such as sRNAs. We found many parallels between the transcriptional progression schemes of LUZ7 and coliphage N4, however, the middle promoter motifs identified in this study, together with the Drc-mediated recruitment of RNAPII (22), hint to different transcriptional initiation mechanisms from middle promoters of the two viruses. Furthermore, our analysis revealed, extensive and time-dependent read-through at phage terminator regions, leading to numerous extended transcription unit variants that incorporate additional genes. Consistent with previous observations in bacteria (15, 69), our results suggest that weak termination sites have an important regulatory role in the transcriptional strategy of phages, allowing them to adjust their dense transcriptional patterns in response to different infection stages. The systematic discovery of dynamic transcription terminators by ONT-cappable-seq could help identify and dissect novel regulatory mechanisms to modulate the strength of terminators in phages and their bacterial hosts.

While the underlying mechanisms through which the annotated transcriptional features orchestrate gene regulation in LUZ7 warrant further investigation, this work already demonstrates the breadth of knowledge that can be gained from ONT-cappable-seq over classical RNA-seq approaches, which generally fail to capture primary RNA species and delineate transcriptional boundaries. Although this study primarily focused on viral transcripts, similar analyses can be performed on the bacterial side to shed further light on the regulatory features that govern the complex interaction between phages and their hosts.

In recent years, several full-length RNA-seq protocols have emerged to comprehensively characterize prokaryotic transcripts (14–16). Notably, the SMRT-Cappable-seq protocol includes size-selection steps to eliminate all cDNA fragments below 1 kb from the final library (15). This approach would be less suitable to study the LUZ7 transcriptome, as illustrated by the staggering amount of transcripts encoding small proteins such as gp01 and Drc, and the unique intergenic sRNA transcription hotspot revealed in this work. Nevertheless, future efforts to mitigate the challenges in end-to-end sequencing of particularly long RNA molecules could further enhance the annotation of transcriptional boundaries and operon organizations by ONT-cappable-seq and may increase the number of detected TSS and TTS in this study.

Taken together, this work introduced ONT-cappable-seq as a novel method to profile prokaryotic transcripts in full-length and yielded a first global dataset of TSSs, TTSs and operons in a phage infection model. In particular, we found that phages whose genomes are densely encoded with small genes and that rely on complex transcriptional regulation systems, are a promising sandbox for the application of ONT-cappable-seq. In phage biology, our current understanding of transcriptome architecture and temporal gene regulation is mainly restricted to a handful of model phages. We envision that the pipeline provided in this study can serve as a gateway to move beyond these model phages and chart the transcriptional landscapes of diverse and uncharacterized phages in unprecedented detail. In addition to the PromethION sequencing platform used in this study, our technique is also compatible with the portable MinION device, offering a highly accessible and cost-effective approach to obtain a bird’s eye view of the transcriptional scheme of phages. In the future, the global elucidation of viral transcriptional regulatory sequences can help unveil the complexity of phage-host interactions, which can ultimately drive phage-based applications in medicine and biotechnology forward.

## DATA AVAILABILITY

The resolved genome of *P. aeruginosa* strain US449 was deposited in NCBI GenBank (accession number CP091880). Raw RNA sequencing files are deposited under GEO accession number GSE196845. All scripts and codes used in this study are made available on Github (https://github.com/LoGT-KULeuven/ONT-cappable-seq). Any additional information is accessible from the authors upon request.

## ACKNOWLEDGEMENTS

We thank Genomics Core Leuven en its staff for providing the PromethION flow cell and assistance with sequencing.

## SUPPLEMENTARY FIGURES

**Supplementary Figure S1:**
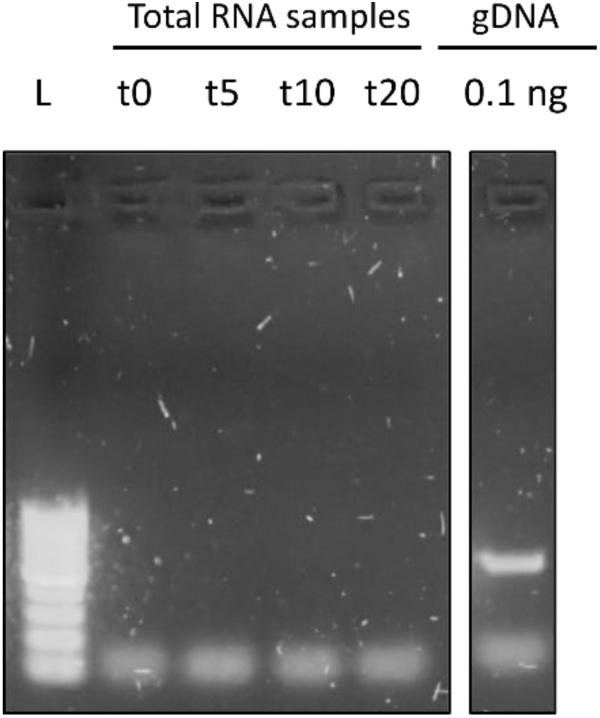
validation of successful DNase treatment of total RNA samples. For all RNA samples (t0, t5, t10 and t20), successful removal of genomic DNA was confirmed by PCR using a host-specific primer pair. As a control, the PCR was performed on a sample containing 0.1 ng of gDNA of strain US449. The PCR products were run on a 1% agarose gel. As a reference, the 100 bp GeneRuler ladder is used (L).

**Supplementary Figure S2:**
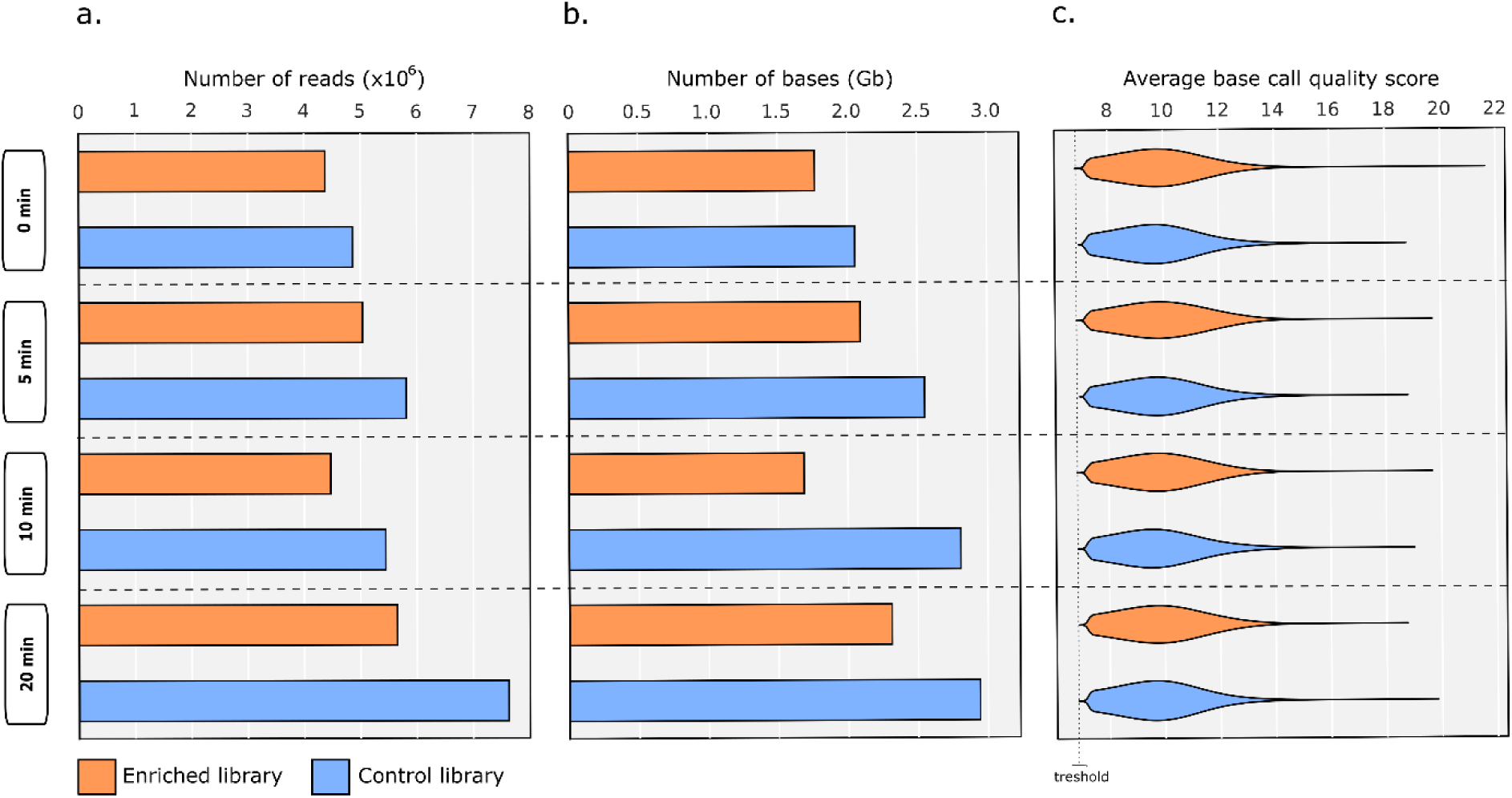
Sequencing throughput and raw read qualities. **a**. Total number of reads for each timepoint after sequencing the enriched library (orange) and the control library (blue). **b**. Total sequencing throughput in gigabases. **c**. distribution of average base call quality score (Phred-like score) of the raw reads.

**Supplementary Figure S3:**
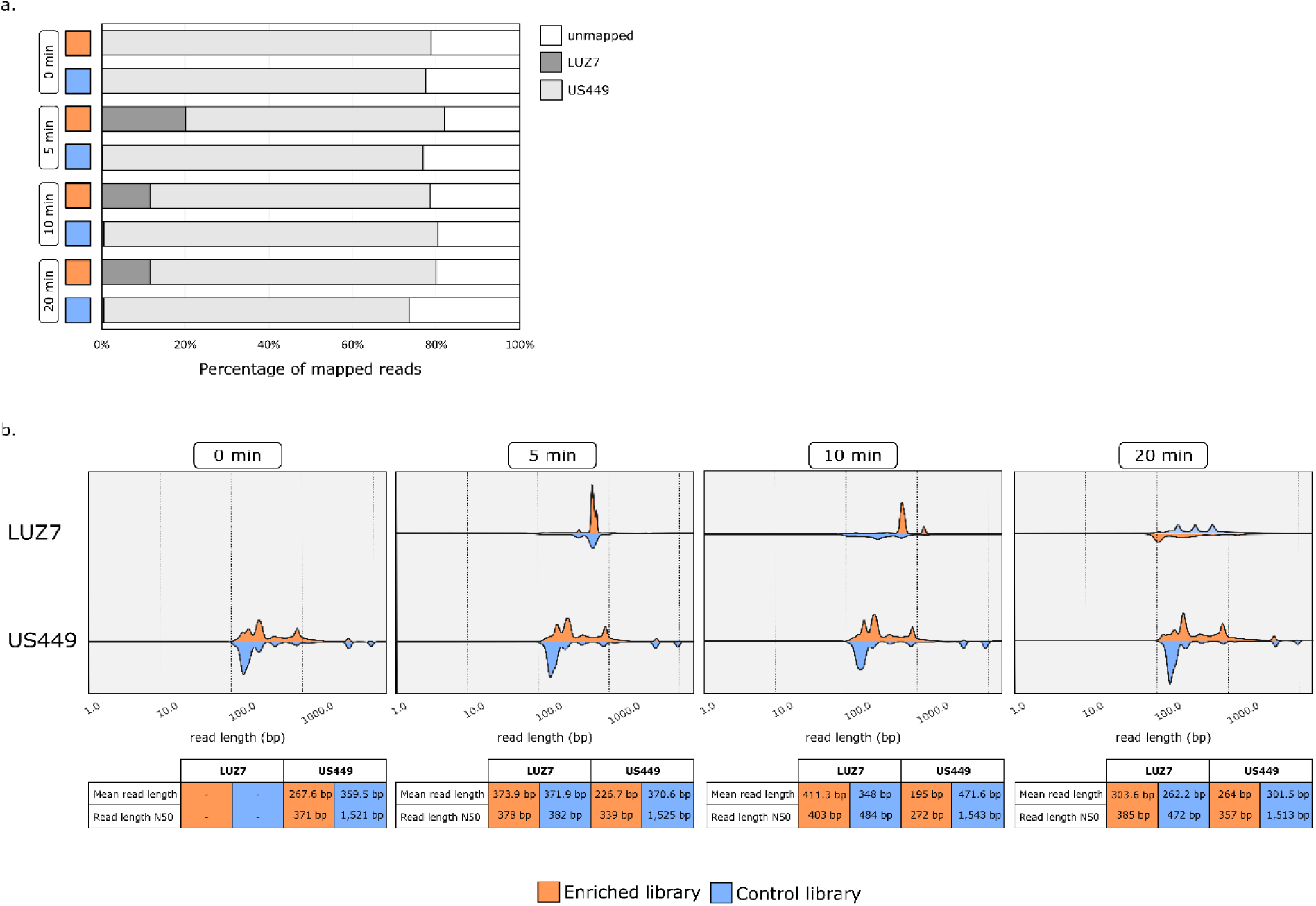
Mapped read characteristics. **a**. Proportion of mapped reads per sample. **b**. Read length distributions of phage-derived and host-derived reads for each infection timepoint for the enriched library (orange) and control (blue).

**Supplementary Figure S4:**
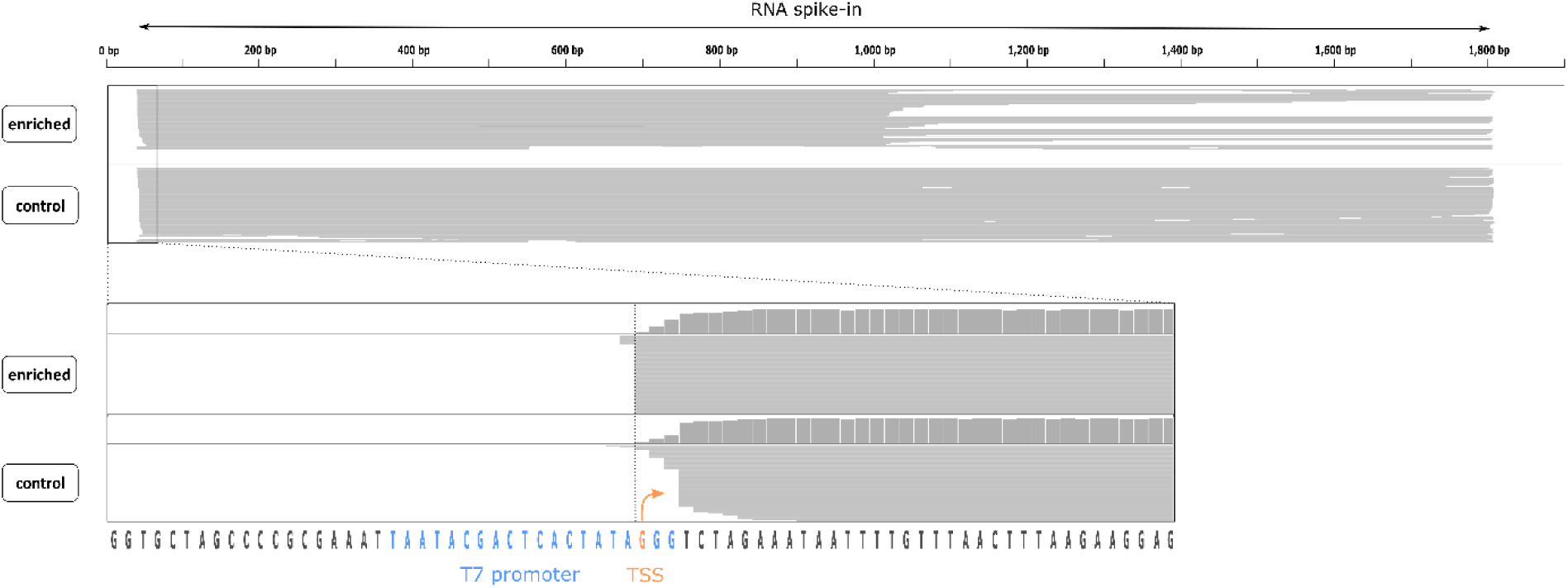
Evaluation of TSS identification of *in vitro* synthesized RNA spike-in by ONT-cappable-seq. Visualization of the IGV tracks of the enriched and control datasets for an *in vitro* transcribed RNA spike-in (1.8kb) under the control of a phage T7 promoter. The TSS identified by ONT-cappable-seq matches the expected TSS of the T7 promoter.

**Supplementary Figure S5:**
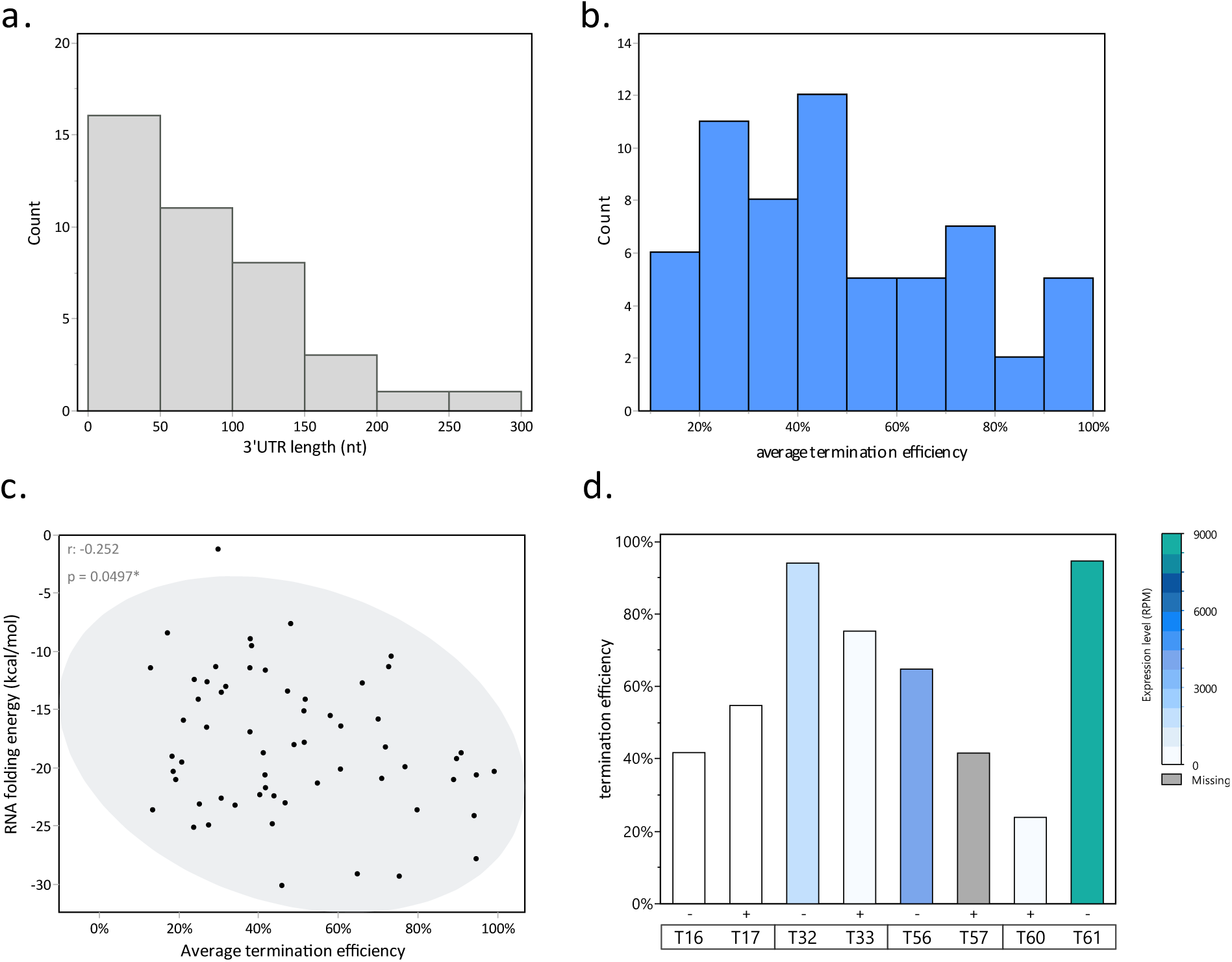
Characteristics of the identified phage terminator regions. **a**. Bar plot showing the distribution of the distance (nt) between the identified TTSs and the stop codon of the nearest annotated gene encoded on the same strand (within 1000 bp), defined as the 3’ UTR. **b**. Distribution of the average transcription termination efficiencies of the terminators found by ONT-cappable-seq. **c**. The RNA folding energy of the terminators as a function of their average termination efficiency. The 95% density ellipse reveals a negative correlation between both parameters (R = -0.267, p < 0.05). **d**. Termination efficiencies of the bidirectional terminator regions in both orientations. For each TTS in the overlapping terminator pair, the colour of the bars indicates the expression level of the gene encoded upstream the TTS. As T57 is localized >1000 nt from the nearest annotated gene with the same orientation, there was no gene expression level associated with T57 (gray bar).

